## Supplementary Figures for "Inverse-folding design of yeast telomerase RNA increases activity *in vitro*"

### Supplemental Figure 1

#### RNA nucleotide sequences of inverse-designed TLC1 alleles

Nucleotide substitution mutations were made in TLC1, without deletions or insertions, in non-essential regions previously deleted in Mini-T (Zappulla *et al.*, *NSMB* 2005) or stiffened in TSA-T (Lebo and Zappulla, *RNA* 2012). Mutated nucleotides are indicated in red. The terminal arm is highlighted in green (with 15 nucleotide substitutions), the Ku arm in orange (with 11–62 substitutions), the Est1 arm in blue (with 33 substitutions), and the template in gray.

(A) TLC1

(B) DA-TLC1

(C) DET-TLC1

(D) DK-TLC1

(E) DPhyK-DA-TLC1

(F) DPhyK-TLC1

#### Supp. Fig. 1A: TLC1

| | | |
|---|---|---|
| 1 | GAGAGGAAGAUAGGUACCCUAUGAAAAUGUCAAUGGCUGUUGCGUUUGCU | Terminal arm (5') |
| 51 | UAAUCGUAUUUUUUUUUUUUUCAGUCCGUGUUUUUUGUACAUUCUACGUU | |
| 101 | UGAGUUUUCCAUCAUGCAGGCCUCAGAAAUUUGGUAGGCACUCGAUGGUG | |
| 151 | AAGAGAUAGUGUCGGAUUUCGGAUUGAUCUUUCAGUUGAUAGCCUGCUGC | |
| 201 | UCUUUUCUUUUCCAAAGAAUUUCGAGUAUGCUGGUGUCAGUGUAGAUGCU | Ku arm |
| 251 | UGUGUGUGCGCAAUUUGUGGUUUUUUAUUGUGUUUCUACUUAUAGAUGGC | |
| 301 | UAAAAUCUGAGUUUAGAAAAUGCAAACCGUAAAUUCUUAAACACUGCUAU | |
| 351 | UGCAUUUAGUUGCUAAAGCAGUGUUUUUGAACUUAUUCCUGUUAUUCCUU | |
| 401 | CUUCGUACCGAUCCUCUUCUCGACCUAACCUUUUAAUUACCAUGGGAAGC | |
| 451 | CUACCAUCACCACACCCACACACAAAUGUUACAGCUAAUUGUUUAUUAGC | template |
| 501 | AAAGUUUGCACGAGUUCGCUGUUUAUUUUUUUCUCGUUUUCUUAUACCUA | |
| 551 | GUAUUUUUUCUGACACUGUUUAAGGUGACAGAAAAAAAGGAGUUUAAGUU | Est1 arm |
| 601 | AGAUUUGCAAACAGACGGUGCUAAGCGCUGUCACUUUAUGUCUAUCUUAU | |
| 651 | CGUUAACUCUGGAAAAAGAAAAAGGAAAAAGAACGUCAGGGAACAUGAGU | |
| 701 | AUAUAUAGAAAUGGUUUAUUCUAGUUUUUUCCGUUUUUUCAGUAGAUUUU | |
| 751 | UGCCUUUAAAAGAAUAAAUCCCACUACAAAAAGGUAAAAUAAAAAAUCUA | |
| 801 | UUCACUGAACUUACUGAUGAAAUUUCCAAAUGUGCCCCGUACAUCGAACG | |
| 851 | AUGUGACAGAGAAAAAUACGAGUAGGUAAAUAAGCCAAAAGGCAAGGGUG | |
| 901 | UCCUUUCUUAAGCAUCGGUUAGGUUUGCGGGCGAUCAGUAACUGAACAAU | |
| 951 | GACACAAGAUCAAGAACGUAAUUUGAGAUUUUUCAAGAUGGUUUUUUUAG | Terminal arm (3') |
| 1001 | GUAUCUAUUAAAACUACUUUGAUGAUCAAUACGGUAUUUUUGUCGCAUUA | |
| 1051 | UUUUCCAAGCGGAAGGAACCGUGUGUUCAUUUUAUGAAUCUUGGUGUUGU | |
| 1101 | AUUCACAGCUACUUCUCCUAAUGCCUUCGAUGCAUUUAGAUAAUUUUUGG | |
| 1151 | AAACAUU | |

### Supp. Fig. 1B: DA-TLC1

1 GAGAGGAAGAUAGGUACCCUAUGAAAAUGCCAAUGGCUGUUGCGUUUGCC
51 UAAUCGUAUUUUUCUCUUUUUCAGUCCGUGUUCUUUGUACAUUCCACGUU
101 UGAGUGUGCCAUCAUGCAGGCCUCAGAAAUUUGGUAGGCACCCGAUGGUG
151 AAGAGACAGUGUCGGCUUUCGGACCGAUCUUUCAGUUGAUAGCCUGCUGC
201 UCUUUUCUUUUCCAAAGAAUUUCGAGUAUGCUGGUGUCAGUGUAGAUGCU
251 UGUGUGUGCGCAAUUUGUGGUUUUUUAUUGUGUUUCUACUUAUAGAUGGC
301 UAAAAUCUGAGUUUAGAAAAUGCAAACCGUAAAUUCUUAAACACUGCUAU
351 UGCAUUUAGUUGCUAAAGCAGUGUUUUUGAACUUAUUCCUGUUAUUCCUU
401 CUUCGUACCGGUCCUUUGCUCGACCUGACCUCUUAAUCACCAUGGGAAGC
451 CUACCAUCACCACACCCACACACAAAUGUUACAGCUAAUUGUUUAUUAGC
501 AAAGUUUGGUCGACUCCUCUGUUUAUCGUGUUCCAAGUGUCCUUUUCCUA
551 GUAUUUUUUCUGACACUGUUUAAGGUGACAGAAAAAAAGGAGUUUAAGUU
601 AGAUUUGCAAACAGACGGUGCUAAGCGCUGUCACUUUAUGUCUAUCUUAU
651 CGUUAACUCUGGUUUAGGACACAGGAACACGACCUUCAGUGGAGACGUCC
701 AUAUAUAGAAAUGGUUUAUUCUAGUUUUUUCCGUUUUUUCAGUAGAUUUU
751 UGCCUUUAAAAGAAUAAAUCCCACUACAAAAAGGUAAAAUAAAAAAUCUA
801 UUCACUGAACUUACUGAUGCACUCUCCAAACGUGCCCCGUACAUCGAACG
851 ACGUGACAGAGAAAAAUACGAGUAGGCAAACAAGCCAAAAGGCAAGGGUG
901 UCCUUUCUUAAGCAUCGGUUAGGUUUGCGGGCGAUCAGUAACUGAACAAU
951 GACACAAGAUCAAGAACGUAAUUUGAGAUUUUUCAAGAUGGUUUUUUUAG
1001 GUAUCUAUUAAAACUACUUUGAUGAUCAAUACGGUAUUUUUGUCGCAUUA
1051 UUUUCCAAGCGGAAGGAACCGUGUGUUCAUUUUAUGAAUCUUGGUGUUGU
1101 AUUCACAGCUACUUCUCCUAAUGCCUUCGAUGCAUUUAGAUAAUUUUUGG
1151 AAACAUU

Terminal arm (5')
Ku arm
template
Est1 arm
Terminal arm (3')

### Supp. Fig. 1C: DET-TLC1

1 GAGAGGAAGAUAGGUACCCUAUGAAAAUGCCAAUGGCUGUUGCGUUUGCC
51 UAAUCGUAUUUUUCUCUUUUUCAGUCCGUGUUCUUUGUACAUUCCACGUU

Terminal arm (5')

101 UGAGUGUGCCAUCAUGCAGGCCUCAGAAAUUUGGUAGGCACUCGAUGGUG
151 AAGAGAUAGUGUCGGAUUUCGGAUUGAUCUUUCAGUUGAUAGCCUGCUGC
201 UCUUUUCUUUUCCAAAGAAUUUCGAGUAUGCUGGUGUCAGUGUAGAUGCU
251 UGUGUGUGCGCAAUUUGUGGUUUUUUAUUGUGUUUCUACUUAUAGAUGGC
301 UAAAAUCUGAGUUUAGAAAAUGCAAACCGUAAAUUCUUAAACACUGCUAU
351 UGCAUUUAGUUGCUAAAGCAGUGUUUUUGAACUUAUUCCUGUUAUUCCUU
401 CUUCGUACCGAUCCUCUUCUCGACCUAACCUUUUAAUUACCAUGGGAAGC

Ku arm

451 CUACCAUCACCACACCCACACACAAAUGUUACAGCUAAUUGUUUAUUAGC

template

501 AAAGUUUGGUCGACUCCUCUGUUUAUCGUGUUCCAAGUGUCCUUUUCCUA
551 GUAUUUUUUCUGACACUGUUUAAGGUGACAGAAAAAAAGGAGUUUAAGUU
601 AGAUUUGCAAACAGACGGUGCUAAGCGCUGUCACUUUAUGUCUAUCUUAU
651 CGUUAACUCUGGUUUAGGACACAGGAACACGACCUUCAGUGGAGACGUCC

Est1 arm

701 AUAUAUAGAAAUGGUUUAUUCUAGUUUUUUCCGUUUUUUCAGUAGAUUUU
751 UGCCUUUAAAAGAAUAAAUCCCACUACAAAAAGGUAAAAUAAAAAAUCUA
801 UUCACUGAACUUACUGAUGCACUCUCCAAACGUGCCCCGUACAUCGAACG
851 ACGUGACAGAGAAAAAUACGAGUAGGCAAACAAGCCAAAAGGCAAGGGUG
901 UCCUUUCUUAAGCAUCGGUUAGGUUUGCGGGCGAUCAGUAACUGAACAAU
951 GACACAAGAUCAAGAACGUAAUUUGAGAUUUUUCAAGAUGGUUUUUUUAG
1001 GUAUCUAUUAAAACUACUUUGAUGAUCAAUACGGUAUUUUUGUCGCAUUA
1051 UUUUCCAAGCGGAAGGAACCGUGUGUUCAUUUUAUGAAUCUUGGUGUUGU
1101 AUUCACAGCUACUUCUCCUAAUGCCUUCGAUGCAUUUAGAUAAUUUUUGG
1151 AAACAUU

Terminal arm (3')

#### Supp. Fig. 1D: DK-TLC1

1 GAGAGGAAGAUAGGUACCCUAUGAAAAUGUCAAUGGCUGUUGCGUUUGCU Terminal arm (5')
51 UAAUCGUAUUUUUUUUUUUUUCAGUCCGUGUUUUUUGUACAUUCUACGUU
101 UGAGUUUUCCAUCAUGCAGGCCUCAGAAAUUUGGUAGGCACCCGAUGGUG
151 AAGAGACAGUGUCGGCUUUCGGACCGAUCUUUCAGUUGAUAGCCUGCUGC
201 UCUUUUCUUUUCCAAAGAAUUUCGAGUAUGCUGGUGUCAGUGUAGAUGCU Ku arm
251 UGUGUGUGCGCAAUUUGUGGUUUUUUAUUGUGUUUCUACUUAUAGAUGGC
301 UAAAAUCUGAGUUUAGAAAAUGCAAACCGUAAAUUCUUAAACACUGCUAU
351 UGCAUUUAGUUGCUAAAGCAGUGUUUUUGAACUUAUUCCUGUUAUUCCUU
401 CUUCGUACCGGUCCUUUGCUCGACCUGACCUCUUAAUCACCAUGGGAAGC
451 CUACCAUCACCACACCCACACACAAAUGUUACAGCUAAUUGUUUAUUAGC template
501 AAAGUUUGCACGAGUUCGCUGUUUAUUUUUUUCUCGUUUUCUUAUACCUA
551 GUAUUUUUUCUGACACUGUUUAAGGUGACAGAAAAAAAGGAGUUUAAGUU Est1 arm
601 AGAUUUGCAAACAGACGGUGCUAAGCGCUGUCACUUUAUGUCUAUCUUAU
651 CGUUAACUCUGGAAAAAGAAAAAGGAAAAAGAACGUCAGGGAACAUGAGU
701 AUAUAUAGAAAUGGUUUAUUCUAGUUUUUUCCGUUUUUUCAGUAGAUUUU
751 UGCCUUUAAAAGAAUAAAUCCCACUACAAAAAGGUAAAAUAAAAAAUCUA
801 UUCACUGAACUUACUGAUGAAAUUUCCAAAUGUGCCCCGUACAUCGAACG
851 AUGUGACAGAGAAAAAUACGAGUAGGUAAAUAAGCCAAAAGGCAAGGGUG
901 UCCUUUCUUAAGCAUCGGUUAGGUUUGCGGGCGAUCAGUAACUGAACAAU
951 GACACAAGAUCAAGAACGUAAUUUGAGAUUUUUCAAGAUGGUUUUUUUAG Terminal arm (3')
1001 GUAUCUAUUAAAACUACUUUGAUGAUCAAUACGGUAUUUUUGUCGCAUUA
1051 UUUUCCAAGCGGAAGGAACCGUGUGUUCAUUUUAUGAAUCUUGGUGUUGU
1101 AUUCACAGCUACUUCUCCUAAUGCCUUCGAUGCAUUUAGAUAAUUUUUGG
1151 AAACAUU

#### Supp. Fig. 1E: DPhyK-DA-TLC1

1 GAGAGGAAGAUAGGUACCCUAUGAAAAUGCCAAUGGCUGUUGCGUUUGCC
51 UAAUCGUAUUUUUCUCUUUUUCAGUCCGUGUUCUUUGUACAUUCCACGUU
101 UGAGUGUGCCAUCAUGCAGGCCUCAGAAAUUUGGUAGGCACCCGAUGGUG
151 AAGAGACAGUGUCGGCUUUCGGACCGUUCUUUUCGGUCAGAGCCUGUCGC
201 UCUUCUCUCUUCCAGAGAGUUUCGUGCACGCUGGCGUCAGUGUAGACGCC
251 UGUGCGUGCUCAAUUUGUGGUUUUUUAUUGUGUUUCUACUUAUAGAUGGC
301 UAAAAUCUGAGUUUAGAAAAUGCAAACCGUAAAUUCUUCUUUCUUCUUCU
351 UGCGUUCAGUUGCUCAACCAGUGUUUUUCCGGUUAUUCCUCCGUUUCUUU
401 CUUUCUUUCGGUCCUUUGCUCGACCUGACCUCUUAAUCACCAUGGGAAGC
451 CUACCAUCACCACACCCACACACAAAUGUUACAGCUAAUUGUUUAUUAGC
501 AAAGUUUGGUCGACUCCUCUGUUUAUCGUGUUCCAAGUGUCCUUUUCCUA
551 GUAUUUUUUCUGACACUGUUUAAGGUGACAGAAAAAAAGGAGUUUAAGUU
601 AGAUUUGCAAACAGACGGUGCUAAGCGCUGUCACUUUAUGUCUAUCUUAU
651 CGUUAACUCUGGUUUAGGACACAGGAACACGACCUUCAGUGGAGACGUCC
701 AUAUAUAGAAAUGGUUUAUUCUAGUUUUUUCCGUUUUUUCAGUAGAUUUU
751 UGCCUUUAAAAGAAUAAAUCCCACUACAAAAAGGUAAAAUAAAAAAUCUA
801 UUCACUGAACUUACUGAUGCACUCUCCAAACGUGCCCCGUACAUCGAACG
851 ACGUGACAGAGAAAAAUACGAGUAGGCAAACAAGCCAAAAGGCAAGGGUG
901 UCCUUUCUUAAGCAUCGGUUAGGUUUGCGGGCGAUCAGUAACUGAACAAU
951 GACACAAGAUCAAGAACGUAAUUUGAGAUUUUUCAAGAUGGUUUUUUUAG
1001 GUAUCUAUUAAAACUACUUUGAUGAUCAAUACGGUAUUUUUGUCGCAUUA
1051 UUUUCCAAGCGGAAGGAACCGUGUGUUCAUUUUAUGAAUCUUGGUGUUGU
1101 AUUCACAGCUACUUCUCCUAAUGCCUUCGAUGCAUUUAGAUAAUUUUUGG
1151 AAACAUU

Terminal arm (5')

Ku arm

template

Est1 arm

Terminal arm (3')

### Supp. Fig. 1F: DPhyK-TLC1

Terminal arm (5')

1 GAGAGGAAGAUAGGUACCCUAUGAAAAUGUCAAUGGCUGUUGCGUUUGCU

51 UAAUCGUAUUUUUUUUUUUUUCAGUCCGUGUUUUUGUACAUUCUACGUU

101 UGAGUUUUCCAUCAUGCAGGCCUCAGAAAUUUGGUAGGCACCCGAUGGUG

Ku arm

151 AAGAGACAGUGUCGGCUUUCGGACCGUUCUUUUCGGUCAGAGCCUGUCGC

201 UCUUCUCUCUUCCAGAGAGUUUCGUGCACGCUGGCGUCAGUGUAGACGCC

251 UGUGCGUGCUCAAUUUGUGGUUUUUUAUUGUGUUUCUACUUAUAGAUGGC

301 UAAAAUCUGAGUUUAGAAAAUGCAAACCGUAAAUUCUUCUUUCUUCUUCU

351 UGCGUUCAGUUGCUCAACCAGUGUUUUUCCGGUUAUUCCUCCGUUUCUUU

401 CUUUCUUUCGGUCCUUUGCUCGACCUGACCUCUUAAUCACCAUGGGAAGC

template

451 CUACCAUCACCACACCCACACACAAAUGUUACAGCUAAUUGUUUAUUAGC

Est1 arm

501 AAAGUUUGCACGAGUUCGCUGUUUAUUUUUUUCUCGUUUUCUUAUACCUA

551 GUAUUUUUUCUGACACUGUUUAAGGUGACAGAAAAAAAGGAGUUUAAGUU

601 AGAUUUGCAAACAGACGGUGCUAAGCGCUGUCACUUUAUGUCUAUCUUAU

651 CGUUAACUCUGGAAAAAGAAAAAGGAAAAAGAACGUCAGGGAACAUGAGU

701 AUAUAUAGAAAUGGUUUAUUCUAGUUUUUUCCGUUUUUUCAGUAGAUUUU

751 UGCCUUUAAAAGAAUAAAUCCCACUACAAAAAGGUAAAAUAAAAAAUCUA

Terminal arm (3')

801 UUCACUGAACUUACUGAUGAAAUUUCCAAAUGUGCCCCGUACAUCGAACG

851 AUGUGACAGAGAAAAAUACGAGUAGGUAAAUAAGCCAAAAGGCAAGGGUG

901 UCCUUUCUUAAGCAUCGGUUAGGUUUGCGGGCGAUCAGUAACUGAACAAU

951 GACACAAGAUCAAGAACGUAAUUUGAGAUUUUUCAAGAUGGUUUUUUUAG

1001 GUAUCUAUUAAAACUACUUUGAUGAUCAAUACGGUAUUUUUGUCGCAUUA

1051 UUUUCCAAGCGGAAGGAACCGUGUGUUCAUUUUAUGAAUCUUGGUGUUGU

1101 AUUCACAGCUACUUCUCCUAAUGCCUUCGAUGCAUUUAGAUAAUUUUUGG

1151 AAACAUU

### Supplemental Figure 2

#### ***Mfold*** **secondary structure predictions of DA-TLC1 alleles**

*Mfold* secondary structure predictions without constraints, with *P-num* output.

(A) TLC1

(B) DA-TLC1

(C) DET-TLC1

(D) DK-TLC1

(E) DPhyK-DA-TLC1

(F) DPhyK-TLC1

### Supp. Fig. 2A: TLC1

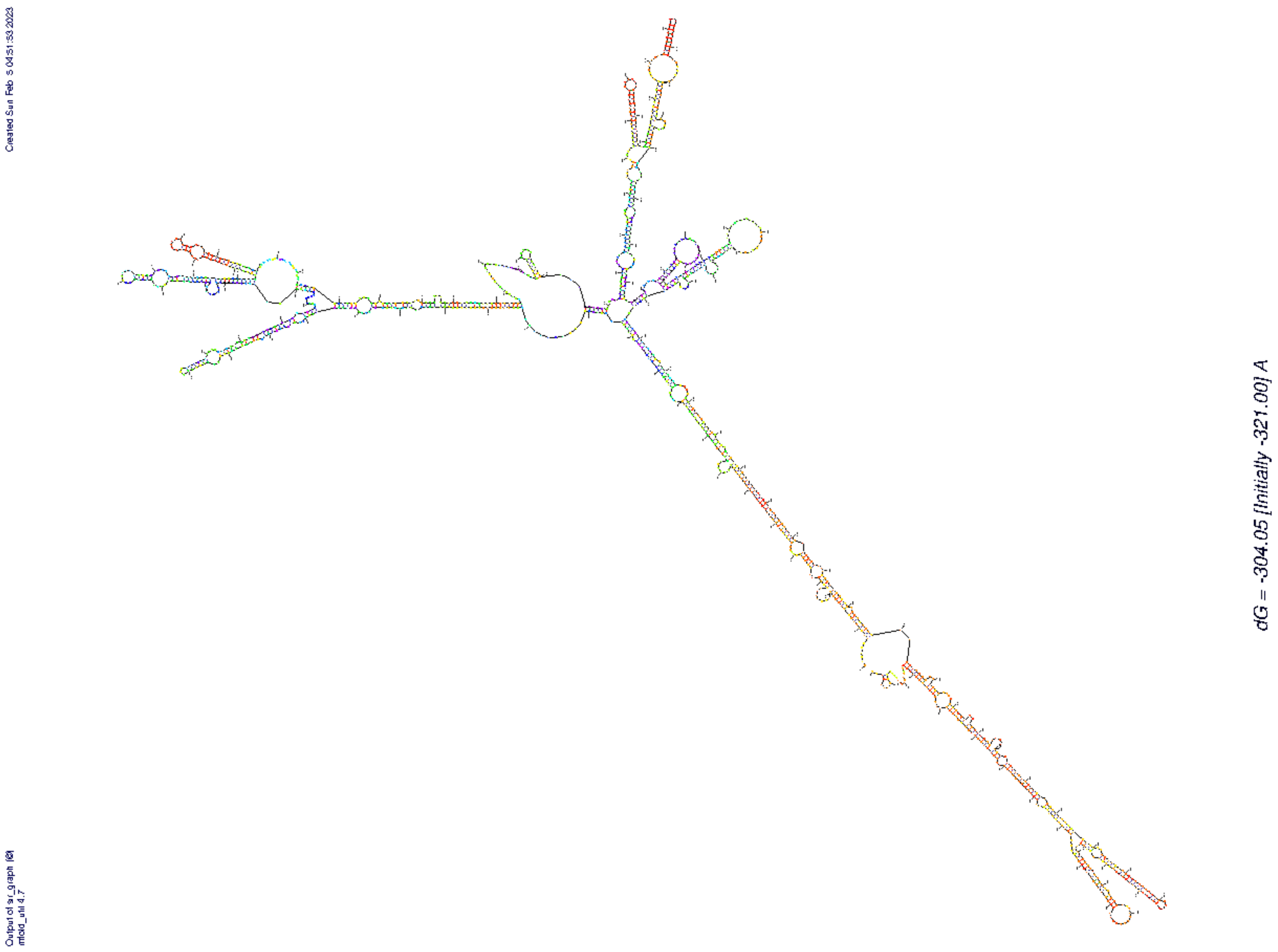

### Supp. Fig. 2B: DA-TLC1

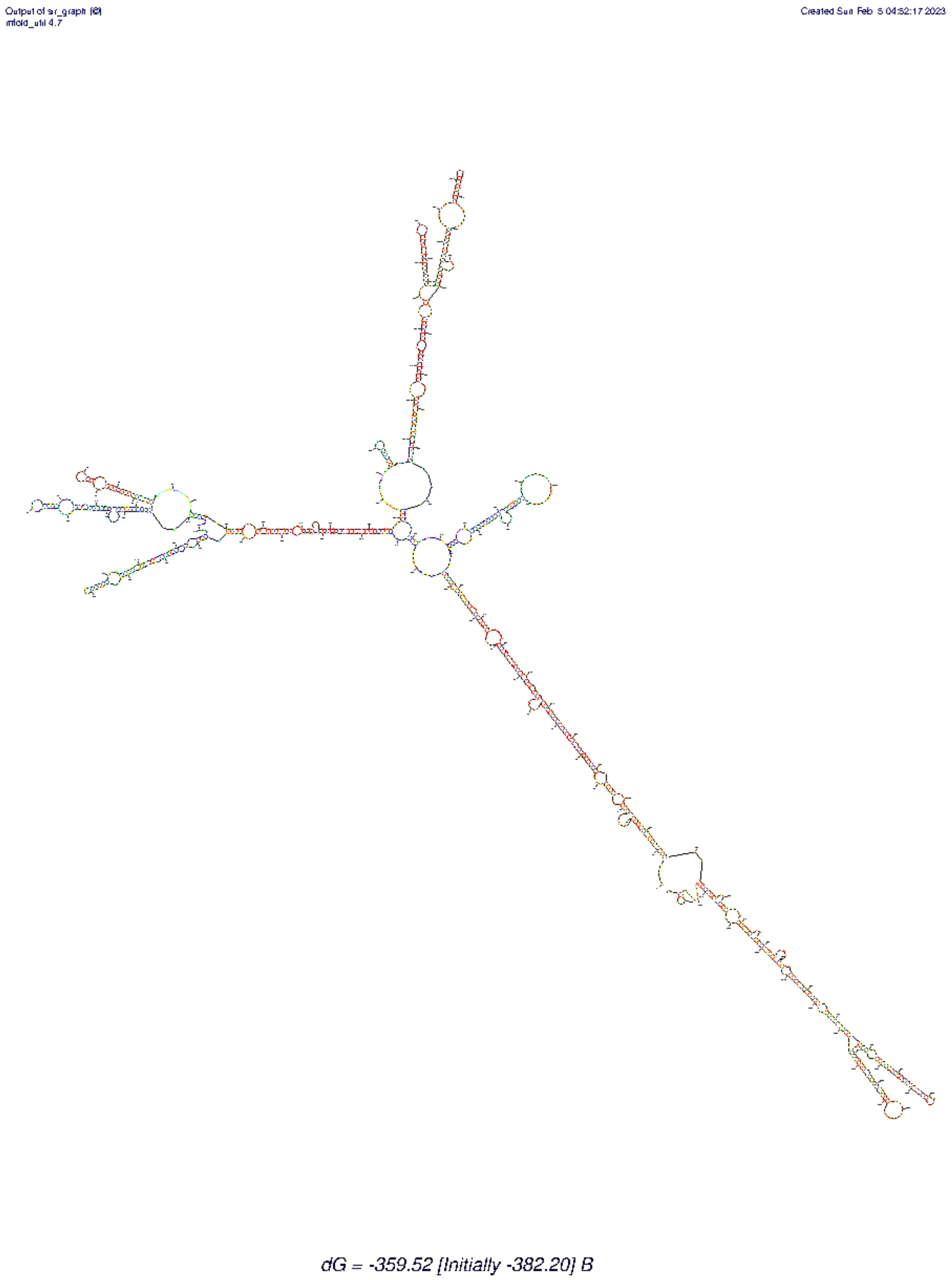

### Supp. Fig. 2C: DET-TLC1

Output of sir_graph (©)
mfold_util 4.7

Created Sun Feb 5 04:54:27 2023

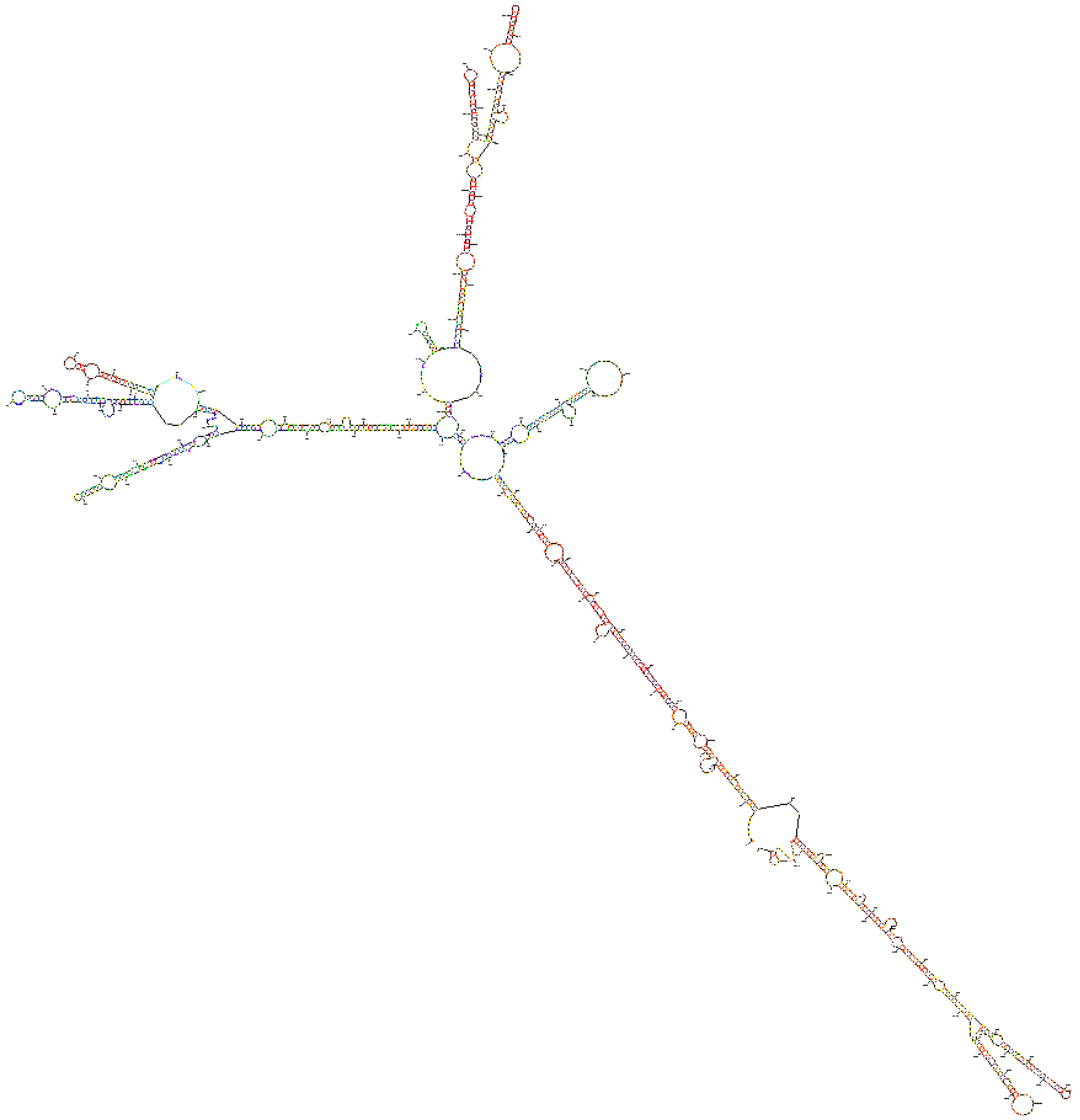

dG = -346.96 [Initially -367.30] C

### Supp. Fig. 2D: DK-TLC1

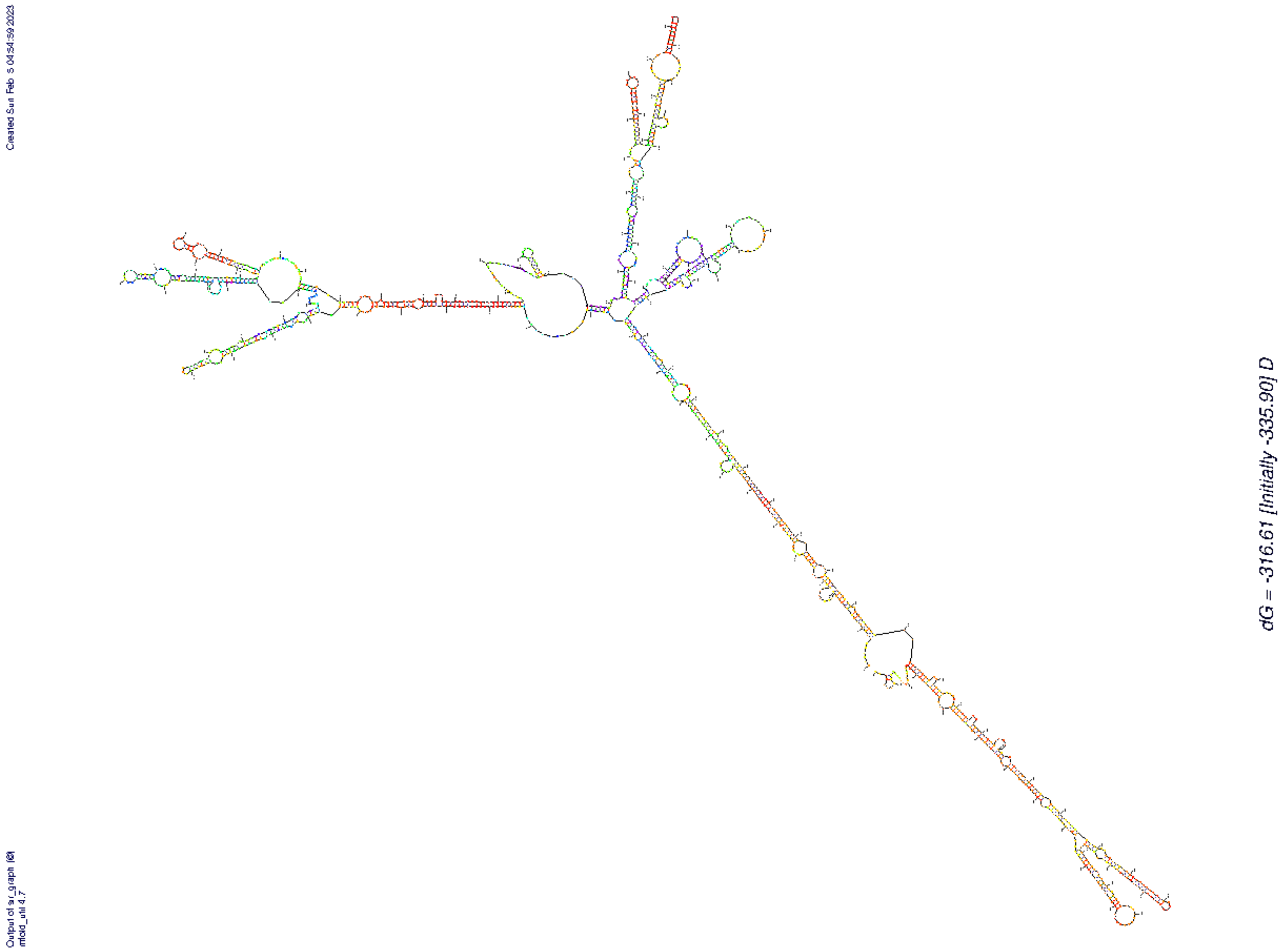

### Supp. Fig. 2E: DPhyK-DA-TLC1

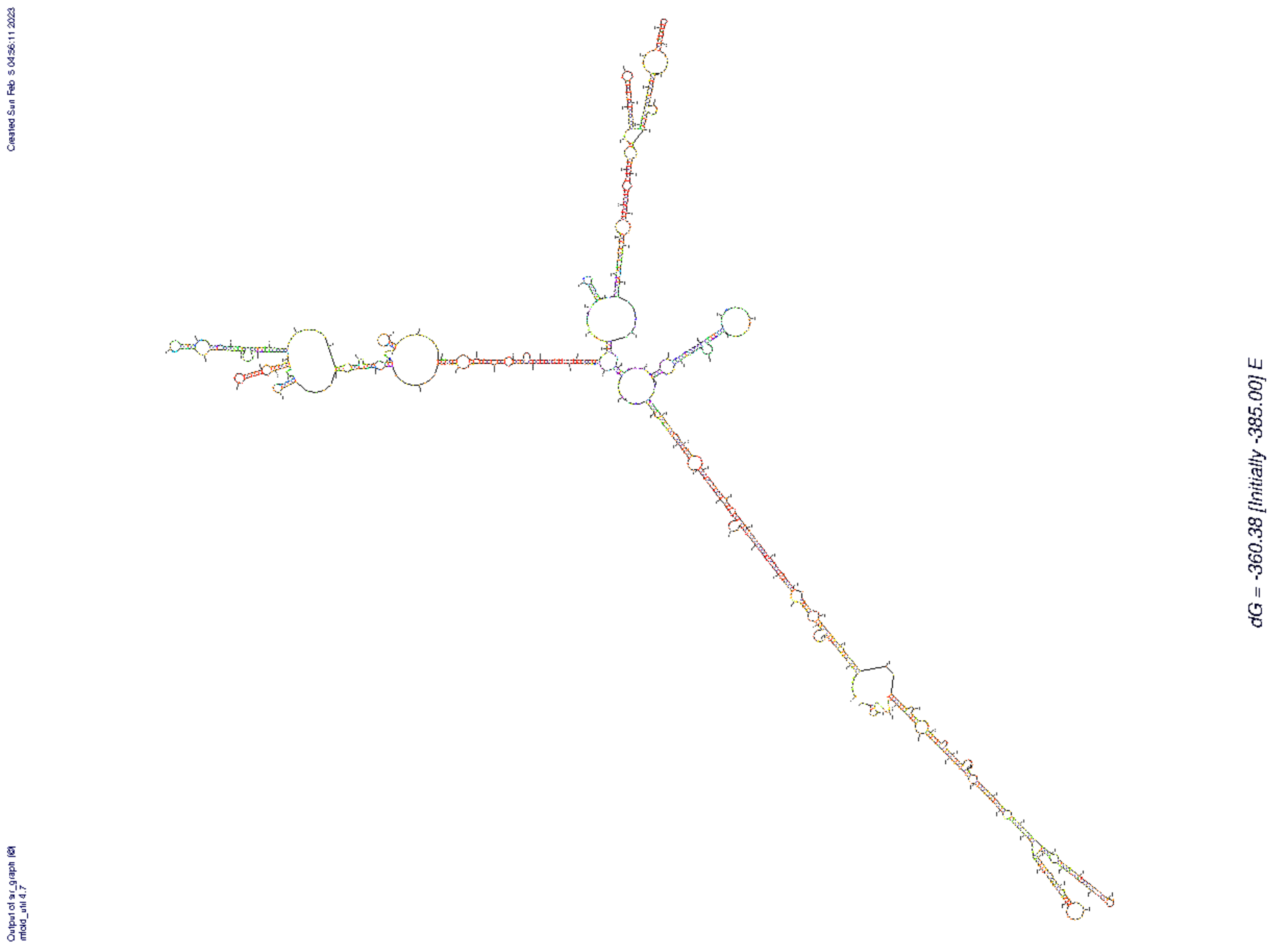

### Supp. Fig. 2F: DPhyK-TLC1

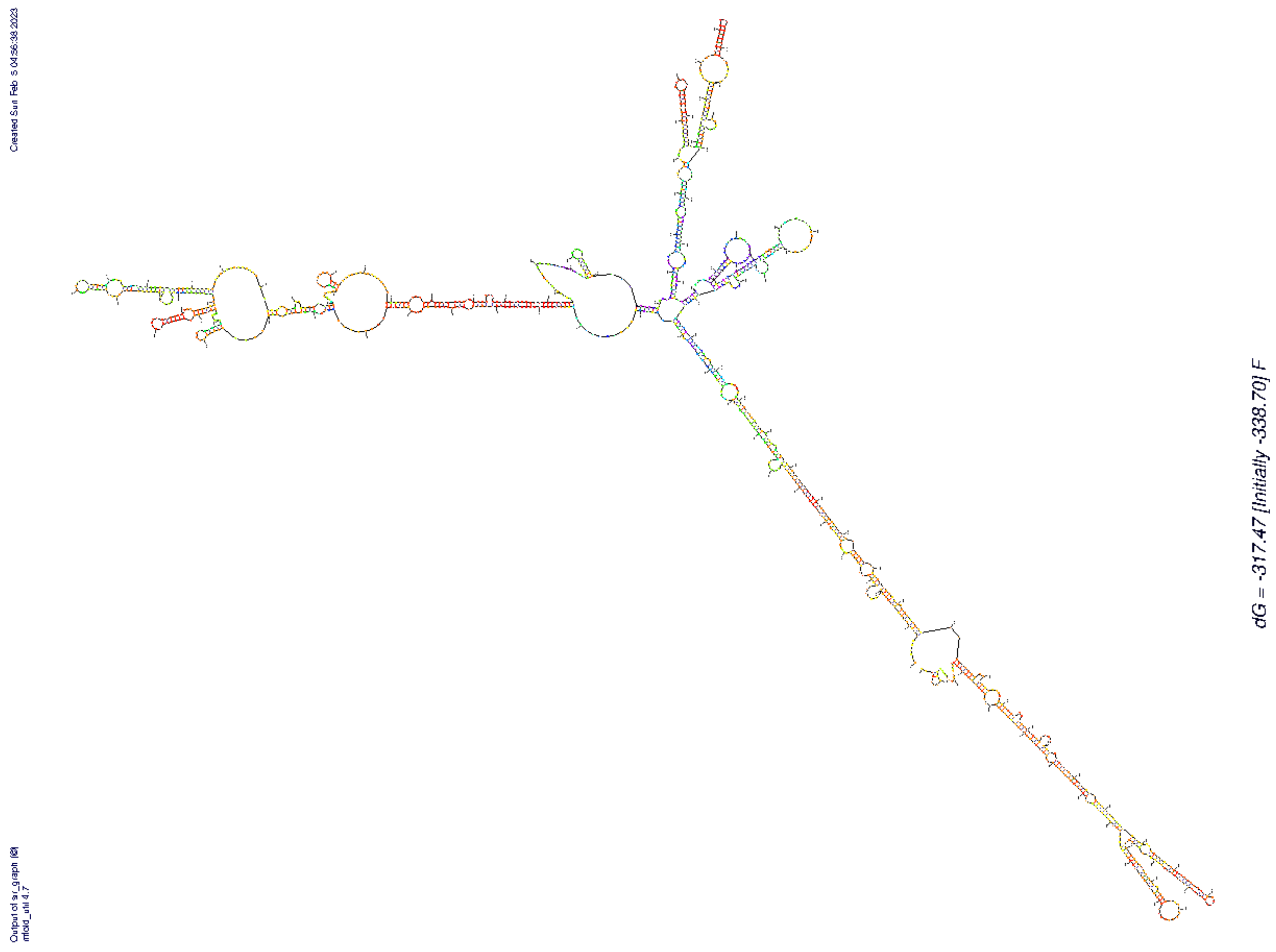

### Supplemental Figure 3

**Comparison of *Mfold* secondary structure predictions for wild-type, inverse-designed, and phylogenetic models of TLC1 arms.** Regions of TLC1 arm secondary structures are shown, comparing the *Mfold* secondary structure prediction and phylogenetically determined (Zappulla and Cech, *PNAS* 2004) structures of wild-type TLC1 to the *Mfold*-predicted structures of the inverse-designed "determined" arms. Nucleotide changes in inverse-designed alleles are circled in red.

(A) Ku-binding arm

(B) Est1-binding arm

(C) Terminal arm

### Supp. Figure 3A

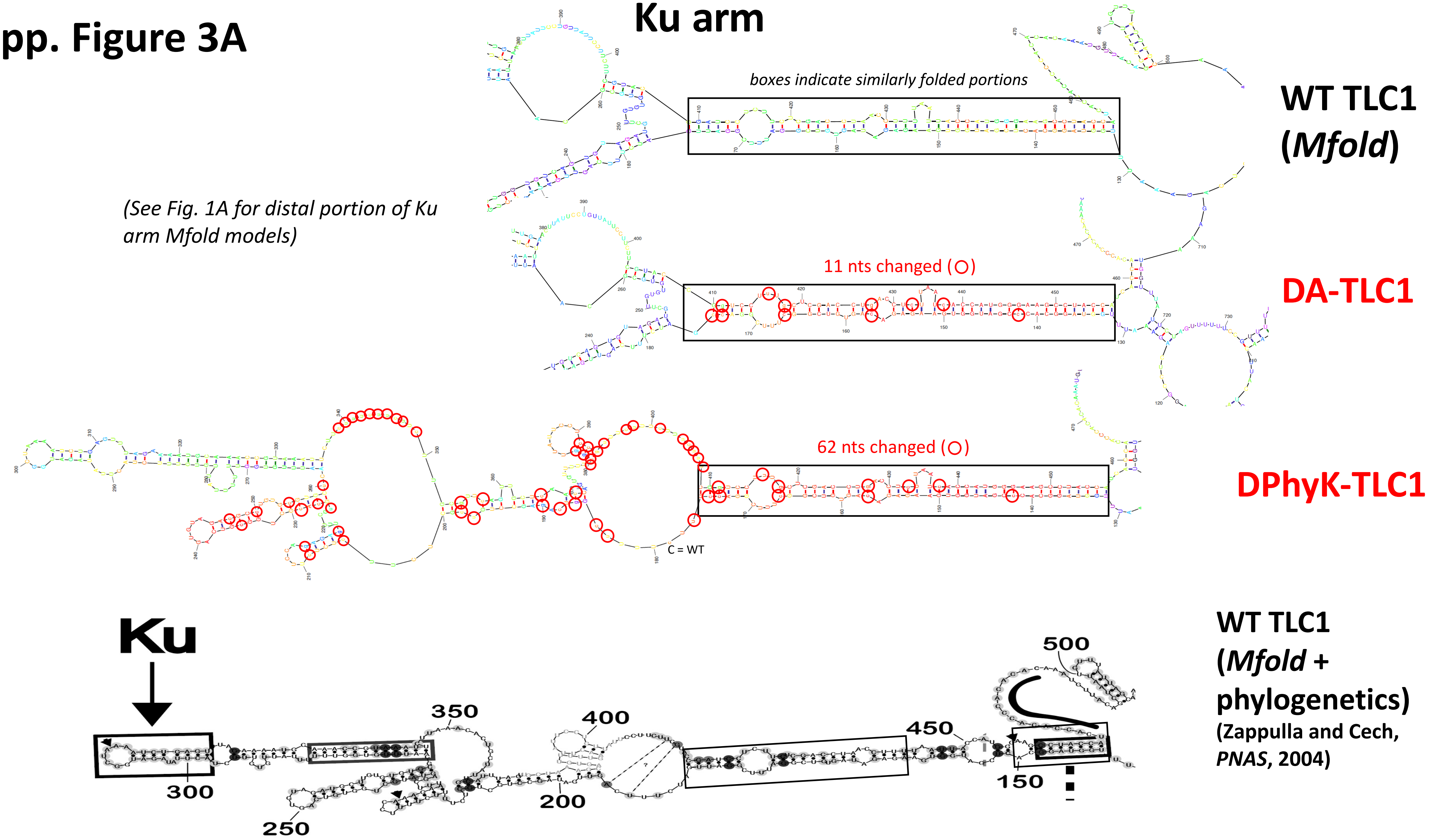

### Supp. Figure 3B

*(See Fig. 1A for distal portion of Est1 arm Mfold models)*

Est1 arm

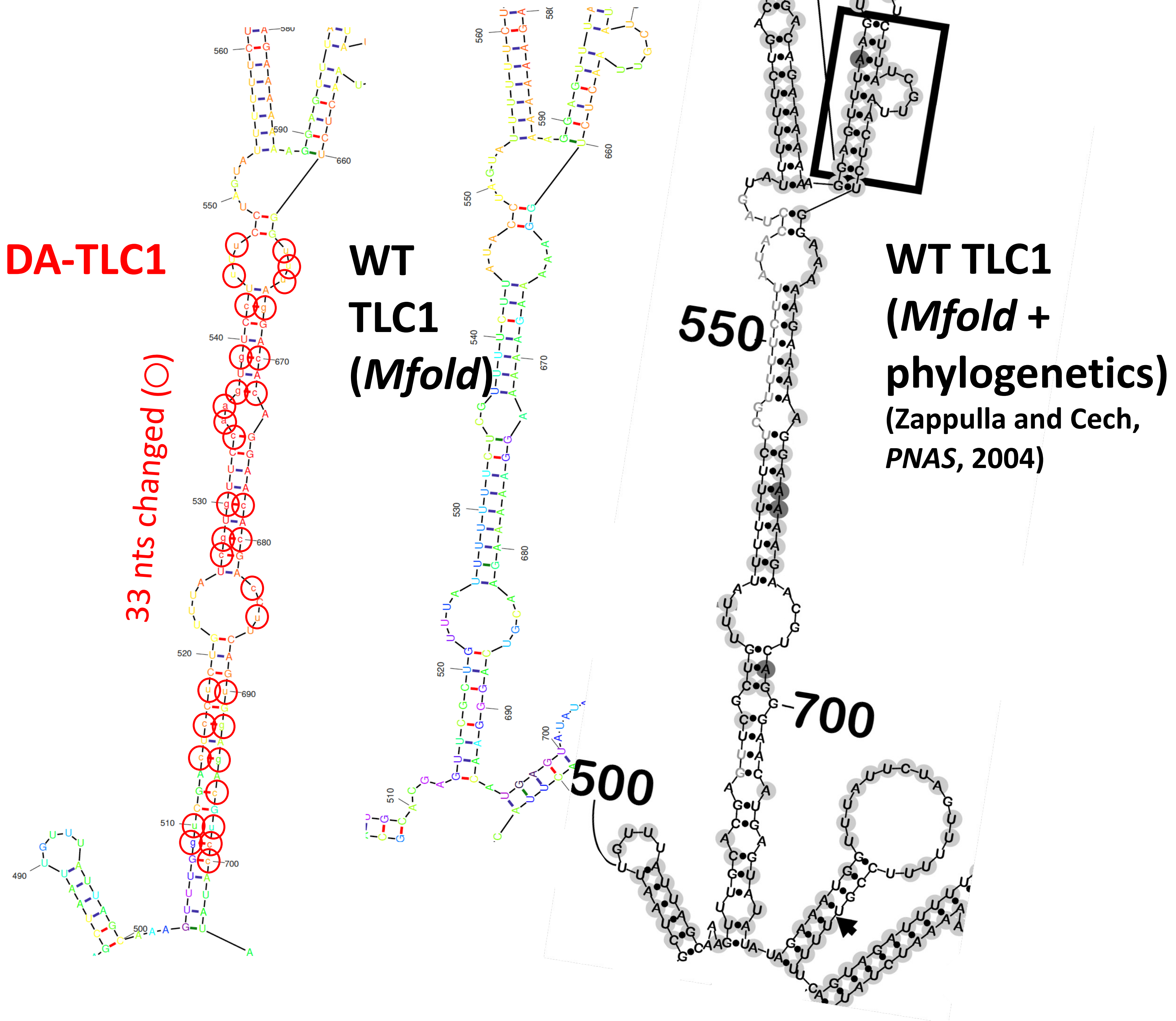

### Supp. Figure 3C

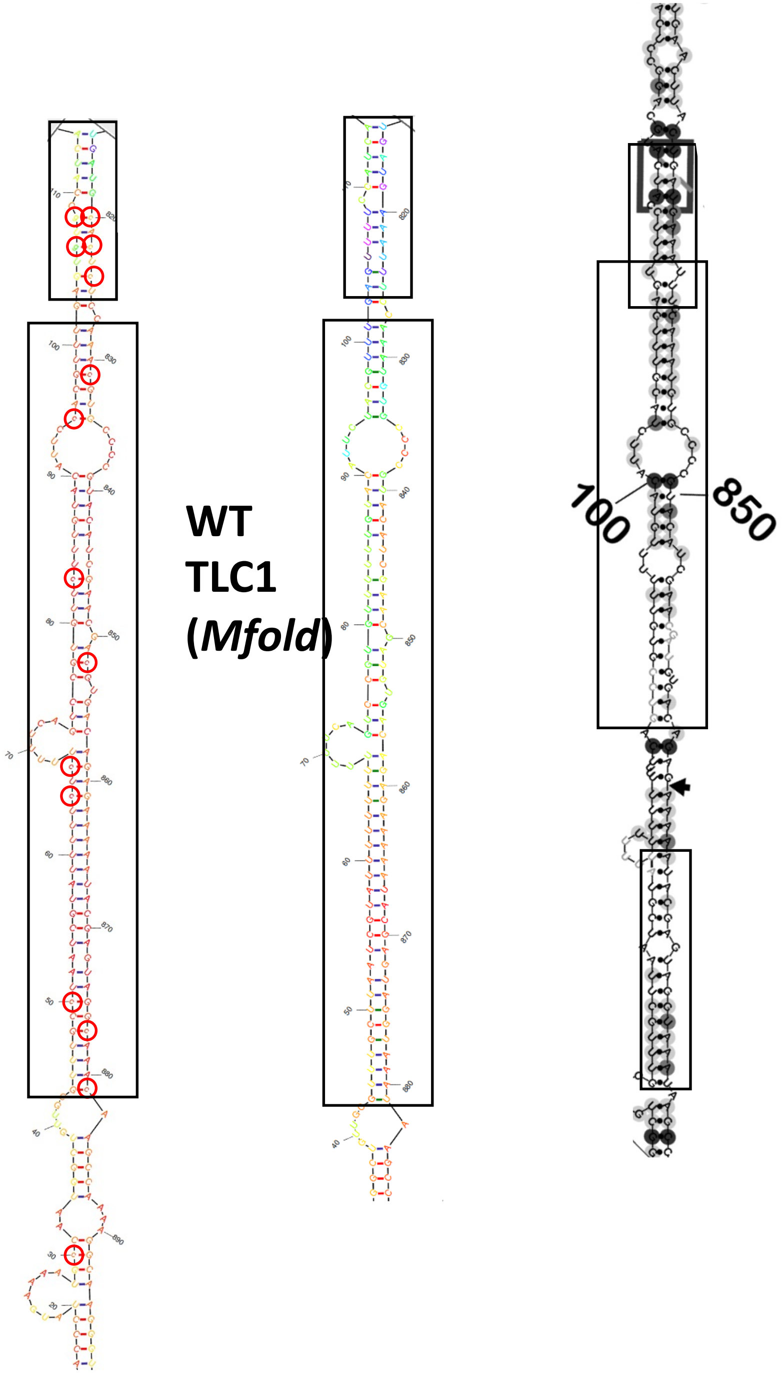


*(See Fig. 1A for distal portion of terminal arm Mfold models)*
